# Inhibition of Alpha Interferon-Induced miR-873 Negatively Affects the Hepatitis B Virus Expression

**DOI:** 10.1101/2024.01.12.575436

**Authors:** Mingsha Zhou, Liu Xing, Jie Fan, Mingming Zhao, Lijuan Chen, Jia Luo, Shan Li, Pan Luo, Yong Duan, Li Zhou

## Abstract

**Background:** Alpha interferon (IFN-α)-based therapy can effectively treat chronic hepatitis B virus (HBV) infection, which remains a serious public health problem. This study aims to identify the potential biomarker of IFN treatment and investigate the effects of miR-873 on chronic hepatitis B (CHB).

**Methods:** We utilized the random forest to screen core differentially expressed miRNAs (DEmiRNAs) based on the miRNA (GSE29911) dataset. DEmiRNAs expressions were determined in clinical samples from CHB patients and normal controls. HBV-transfected hepatoma cell lines were constructed for vitro study. Quantitative real-time reverse transcription PCR (RT-qPCR) was used to determine miRNA mRNA expressions. ELISA assays were used to detect the secretion levels of HBsAg and HBeAg.

**Results:** Boruta feature selection revealed miR-873 hinting at a predictive marker for IFN response. MiR-873 was significantly up-regulated in the CHB patients, as compared with normal controls (*P<*0.001). Increased expression of miR-873 was also found in HBV-infected cells (*P<*0.001). With IFN-α treatment, miR-873 was significantly down-regulated in HBV-infected Huh-7 and HepG-NTCP cells (*P<*0.05). IFN-α treatment and miR-873 inhibitor significantly reduced HBsAg and HBeAg level (*P*<0.05). The inhibitory effect was enhanced by blocking the IFN-α-induced miR-873 downregulation by inhibitor transfection (*P*<0.05). In miR-873 mimic-treated cells, the inhibitory effect was greatly supressed by blocking the IFN-α-induced miR-873 downregulation by mimic transfection (*P*< 0.01).

**Conclusion:** We provide evidence that inhibition of miR-873 by IFN-α enhances the antiviral effects of IFN-α. Our study provides a potential strategy to enhance the anti-HBV effect of IFN-α by inhibiting miR-873 expression.

## 1. Introduction

Chronic hepatitis B (CHB), which is triggered by HBV infection, remains a major public health issue. WHO estimates that 296 million people were living with CHB infection in 2019, with 1.5 million new infections each year[1]. Currently, treatment of CHB consists mainly of pegylated alpha interferon (IFN-α) and nucleotide analogs (NAs)[2]. There is limited performance of NAs in reducing hepatitis B surface antigen (HBsAg), despite the effectiveness in reducing virologic indicators[2–5]. In fact, treatment with pegylated IFN results in the highest rate of off-treatment sustained responses among currently available drugs[6]. It has been shown that the clinical application of IFN results in sustained response in a minority of patients with chronic hepatitis B virus (HBV) infection and has considerable side effects[7].

As a member of the type I interferons, IFN-α plays an important role in host antiviral responses, including protection against HBV infection. Clinical and experimental studies have suggested that IFN-based therapy (IFN-α-2b and pegylated IFN-α-2a or –2b) also has a direct antiviral effect[8–11]. For example, IFN reduced HBV cccDNA through APOBEC3-mediated deamination. However, the IFN response has been reported to be inhibited by HBV viral and host factors[12, 13]. Notably, previous studies have shown that IFN-α/β has a dual function against HBV[14]. Nevertheless, these studies strongly suggest that there is significant potential to modulate the effectiveness of IFN-mediated anti-HBV activities.

MicroRNAs (miRNAs) are a class of small endogenous single-stranded non-coding RNAs that function as key regulators in a variety of biological processes such as differentiation, proliferation, apoptosis, invasion, stress and immunity[15, 16]. Due to the high stability of most miRNAs in the circulation and seemingly tissue-specific expression patterns, serum miRNAs have emerged as new candidate biomarkers for the diagnosis of infectious diseases. Interestingly, several miRNAs have been identified with abnormal expressions in the process of HBV infection, thereby leading to viral replication and pathogenesis[17]. In addition, exosomal miRNAs (miR-194-5p and miR-22-3p) screened by next-generation sequencing were identified to predict HBeAg seroconversion in CHB patients treated with peginterferon (Peg-IFN)[18]. Therefore, it is crucial to explore more potential molecular markers of chronic hepatitis B.

Given the importance of IFN-α in HBV infection, our study analyzed the expression profiles of miRNAs associated with IFN response in CHB patients. We identified four miRNAs associated with IFN response and further explored the effect of miR-873 expression on chronic hepatitis B.

## 2. Materials and methods

### 2.1. Data collection

Matrix file of GSE29911 was originated from the GEO (http://www.ncbi.nlm.nih.gov/geo/) database. Gene annotation was based on the platform GPL10406 (Agilent-021827 Human miRNA Microarray Rel12.0, v3.0, 8×15k array). The GSE29911 dataset consisted of 94 plasma samples from CHB patients on IFN therapy, of which 40 were responders to IFN therapy and 54 were not.

### 2.2. Identification of DEmiRNAs

The matrix file of GSE29911 was obtained to screen the DEmiRNAs. In brief, the genes with |logFC| ≥ 2 and *P* value <0.05 were selected as the DEGs for next analysis. The “limma” package of R (version 3.4.0, https://www.r-project.org/) was applied for volcano map.

### 2.3. Algorithm-based screening of miRNAs

Feature selection of DEmiRNAs was performed using the Boruta package based on the Random forest algorithm. Compared to other selection-by-variable method, the Boruta method had fairly low out-of-bag error rates and computation times. Boruta tested whether the importance of each individual variable was significantly higher than the importance of the random variables by repeatedly fitting a random forest model. It was not until all predictor variables were categorized as either confirmed or rejected at an alpha level of 0.05. In present study, 94 chronic hepatitis B patients receiving IFN treatment were randomly assigned to training and test groups. The training set consisted of 29 IFN treatment responders and 36 IFN treatment non-responders. The validation set included 11 IFN treatment responders and 18 IFN treatment non-responders. The KNN, as a powerful algorithm, depends on neighboring (or similar) patterns relative to the query pattern, and an important challenge is to find the best distance or similarity metric. We used four weighted nearest neighbor method based on KNN algorithm (“triangular”, “epanechnikov”, “inv” methods, and the standard “rectangular”) to obtain a better prediction model.

### 2.4. Human serum samples

A total of 18 healthy controls and 18 CHB patients were recruited from Jiulongpo People’s Hospital and Nan’an People’s Hospital, respectively. The diagnosis of CHB was based on the “Guidelines for prevention and treatment of chronic hepatitis B, China (2015)”[19]. Participants who had coinfection with human immunodeficiency virus (HIV) or hepatitis C virus (HCV), autoimmune hepatitis, drug-induced injury, or alcohol abuse were excluded. Seven milliliters of fasting blood sample was collected from each participant during his/her first admission to the hospital.

### 2.5. Separation of PBMCs

Peripheral blood mononuclear cells (PBMCs) were isolated from fresh heparinized whole blood by two consecutive centrifugation steps (1000 g for 30 min at 4 °C and 2000 g for 15 min at 4 °C, respectively). The supernatant serum was recovered and then stored at − 80 °C.

### 2.6. Cell culture and treatments

The HepG2-NTCP and Huh-7 cells were maintained in DMEM supplemented with 10% FBS and 1% penicillin/streptomycin. For transfection studies, HBV genome containing 1.1 copies, miRNA inhibitor (20 nM), miRNA mimic (20 nM) and miRNA negative control (20 nM) were transfected into cells using Lipofectamine 2000 (Invitrogen, USA) according to the instructions of the reagent. HepG2-NTCP and Huh-7 cells were treated with recombinant human IFN-α injection (1000IU/ml) from Anke Biotechnology (Anhui, China) to simulate IFN treatment in patients. After the transfected cells were cultured in 37 ℃ incubator for 48 hours, the supernatant and cells were extracted.

### 2.7. RNA extraction, reverse transcription and real-time quantitative PCR analysis

Total RNA was isolated from PBMCs, HepG2-NTCP and Huh-7 cells with RNAi plus (TaKaRa, Japan) according to the manufacture’s protocol. The concentration of RNA was measured at OD260/280 by a NanoDrop ND-2000 spectrophotometer (Thermo Scientific, Wilmington, Delaware). Reverse transcription was performed with miRNA P-RT Solution Mix and miRNA P-RT EnzymeMix (Sangon Biotech., China). The reaction system contained 1 μg of total small RNA, 5 μl miRNA P-RT Solution Mix, and 1 μl miRNA P-RT EnzymeMix. SYBR Green qPCR Master Mix Kit (MedChemExpress, China) was used to quantify the levels of miRNAs based on the BioRad^®^96 instrument Real-Time PCR System. U6 was selected as endogenous control for miRNA expression analysis. The cycle threshold (CT value) was defined as the number of cycles required for the fluorescent signal to cross the threshold in quantitative PCR. The primer sequences used in this study were listed in Supplementary Table 1.

### 2.8. Detection of HBsAg, HBeAg

The concentration of HBsAg and HBeAg in the culture supernatant were detected by commercial ELISA kits ( Kehua Biotech., Shanghai, China) according to the manufacturer’s protocols.

### 2.9. Statistical Analysis

The statistical data were collected using the SPSS statistical software (version 22.0) (IBM Corporation, Armonk, NY, USA), and diagrammed by GraphPad Prism 9.3.1. *P*<0.05 was considered to be indicative of statistical significance. All data were presented as mean ± SE through at least three independent experiments.

## 3. Results

### 3.1. Featured DEmiRNAs selection based on the Boruta algorithm

In the results, 18 DEmiRNAs (3 up-regulated and 15 down-regulated) were identified from the GSE29911 dataset. The top 10 DEmiRNAs sorted according to | logFC | were annotated with the volcano plot (Figure 1A). Feature screening result based on the Boruta algorithm was shown in Figure 1B. In order of Z-values, the 4 miRNAs most closely associated with IFN response were miR-873, miR-200a, miR-30b and let-7g. We generated KNN models using 4 weighting methods (“triangularv”, “epanechnikov”, “inv”, and “rectangular”) for optimal classification performance. The optimal value plot showed the highest prediction accuracy when using “inv” (Supplementary Figure 1. The prediction model constructed based on the KNN algorithm.).

**Figure 1.**
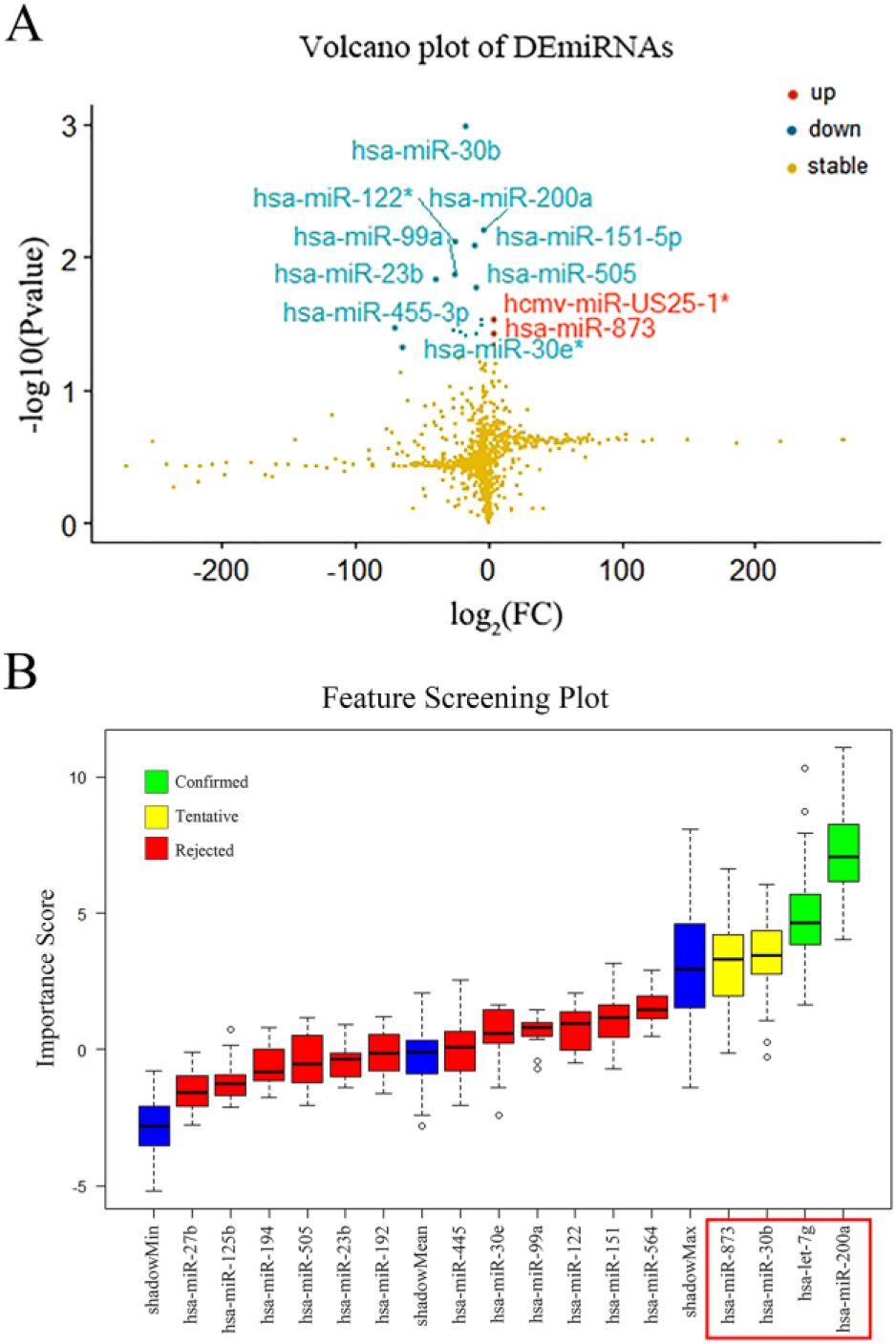
Featured DEmiRNAs selection based on the Boruta algorithm. (A) The volcano plot of 18 DEmiRNAs. Red dots represent the up-regulated genes and green dots represent the down-regulated genes. (B) Feature selection based on the Boruta algorithm. The horizontal axis is the name of each variable, and the vertical axis is the Z-value of each variable. The box plot shows the Z-value of each variable during model calculation. The green boxes represent the comfirmed variables, the yellow represents tentative attributes, and the red represents rejected variables.

### 3.2. Validation of featured DEmiRNAs expression in CHB patients

To understand the expression levels of DEmiRNAs in CHB patients, 4 miRNAs (miR-873, miR-200a, miR-30b, let-7g) were subjected to RT-qPCR analyses. The differences in characteristics between CHB and NC groups are described in Supplementary Table 2. Baseline characteristics analysis of patients with chronic hepatitis B and normal controls. Our results demonstrated that all these DEmiRNAs were up-regulated in CHB group (*P*<0.001), especially miR-873 and let-7g (*P*<0.0001; Figure 2A-D).

**Figure 2.**
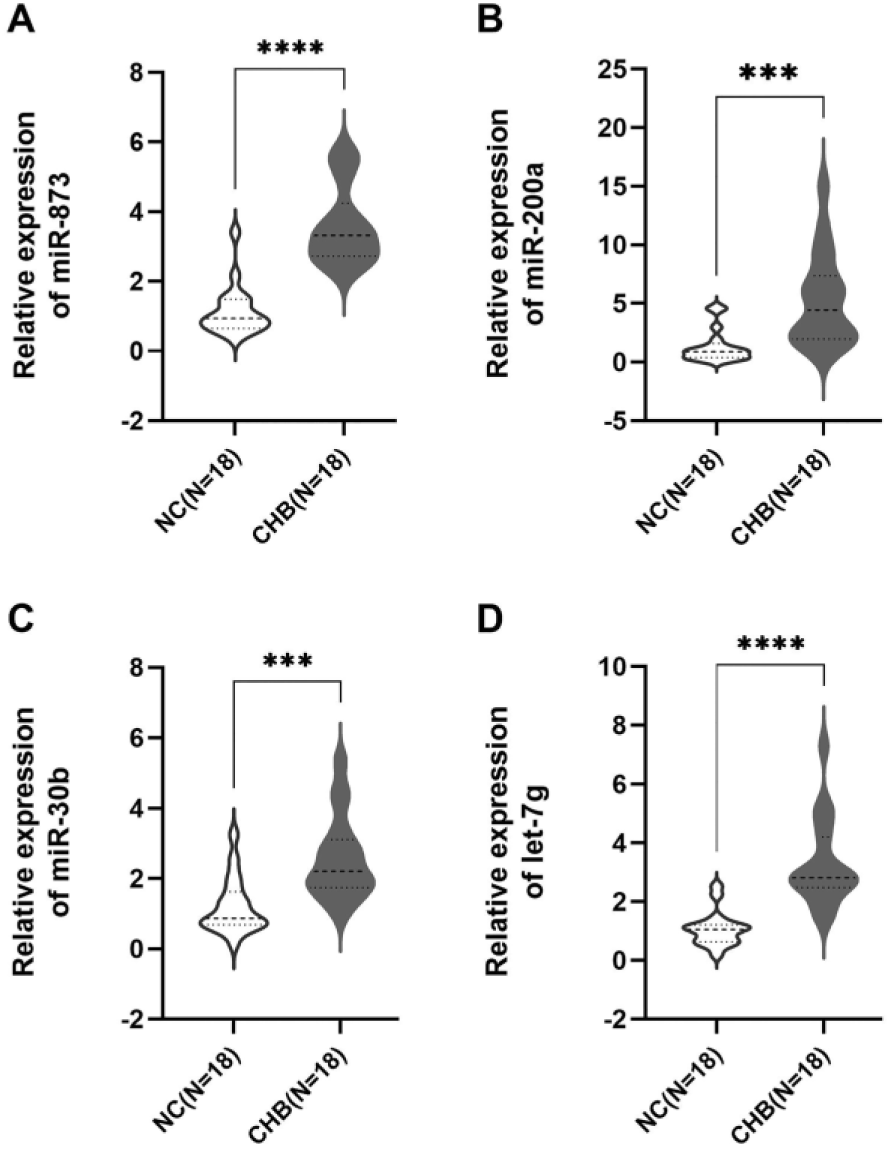
Validation of the relative expression levels of (A) miR-873; (B) miR-100a; (C) miR-30b; (D) let-7g in CHB patients using RT-qPCR. \*\*\**P<*0.001, \*\*\*\**P<*0.0001. Ratios calculated by 2^−ΔΔCt^ method. Data presented as mean ± standard deviation.

### 3.3. Validation of featured DEmiRNAs expression in Huh-7 and HepG2-NTCP cells

In this study, Huh-7 and HepG2-NTCP cells were selected for testing of whether DEmiRNAs was subject to regulation by HBV infection. We analyzed miRNAs expression in HBV plasmid transfection of Huh-7 and HepG2-NTCP cells by RT-qPCR analyses. As shown in Figure 3A, the expression of miR-873 was significantly up-regulated in Huh-7 cells and HepG2-NTCP cells (*P<*0.001). Moreover, our results indicated that miR-200a, miR-30b and let-7g were also upregulated in Huh-7 cells (*P<*0.05) and HepG2-NTCP cells (*P<*0.01), respectively (Figure 3B-D).

**Figure 3.**
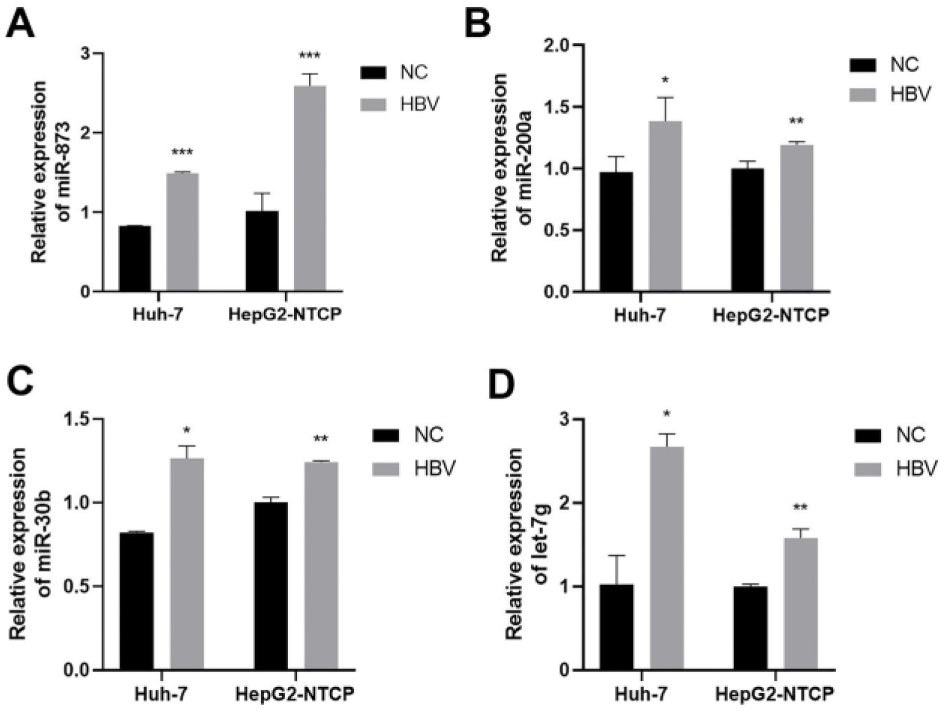
The differential expression of (A) miR-873; (B) miR-200a; (C) miR-30b; (D) let-7g in HBV-infected Huh-7 and HepG2-NTCP cells. \**P<*0.05, \*\**P<*0.01, \*\*\**P<*0.001.

### 3.4. IFN-*α* induces downregulation of miR-873

Huh-7 and HepG2-NTCP cells transfected with the HBV replication plasmid pHBV1.1 were treated with 1,000 U/ml of IFN-α to investigate the expression of miRNAs. The results showed that the expression level of miR-873 was obviously decreased in IFN-treated cells (*P<*0.05, Figure 4A). No significant differences were observed in the levels of miR-200a, miR-30b and let-7g between IFN-treated cells and untreated cells (Figure 4B-D). Therefore, we selected miR-873 as a candidate miRNA for further study. HepG2-NTCP cells were selected for testing of whether miR-873 was subject to regulation by IFN-α because of the knockout inefficiency of miR-873.

**Figure 4.**
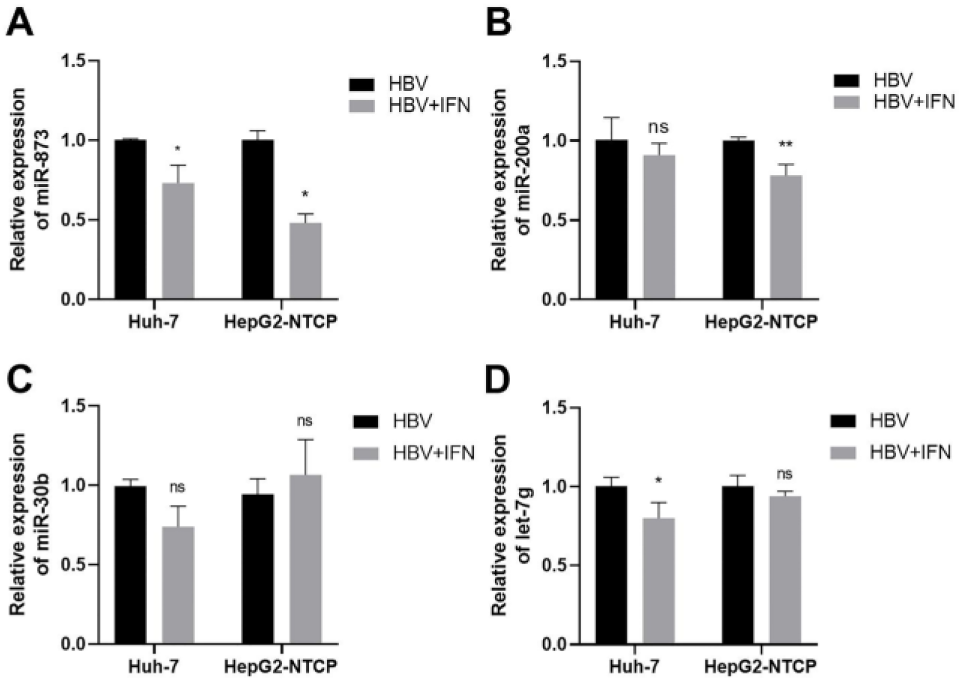
Effect of HBV infection and IFN treatment on (A) miR-873; (B) miR-200a; (C) miR-30b; (D) let-7g expression in Huh-7 and HepG2-NTCP cells. \**P<*0.05, \*\**P<*0.01, ns represents no significance.

### 3.5. IFN-*α* and miR-873 suppresses HBV expression

We analyzed HBV expression in response to IFN-α in HepG2-NTCP cells. As shown in Figure 5A, significant decreases of HBsAg and HBeAg expression were detected in HepG2-NTCP cells 48 h after treatment with IFN-α (*P*<0.05). To investigate the potential effect of IFN-α-induced miR-873 downregulation on HBV expression, HepG2-NTCP cells were treated with 20 nM chemically synthesized miR-873 mimic or 20 nM mimic-NC as a negative control 24 h before transfection with pHBV1.1. The results demonstrated a satisfactory efficiency (*P*<0.01) (Figure 5B). Compared with that in NC-treated cells, significant decreases of HBsAg and HBeAg expression were detected in HepG2-NTCP cells after treatment with miR-873 inhibitor 48 h after transfection (*P*<0.05 for both) (Figure 5C). Further, HepG2-NTCP cells were cotransfected with pHBV1.1 together with 20 nM chemically synthesized miR-873 mimic or 20 nM mimic-NC. Significant increases of HBsAg and HBeAg expression were observed 48 h after miR-873 transfection (*P*<0.01 for both) (Figure 5D).

**Figure 5.**
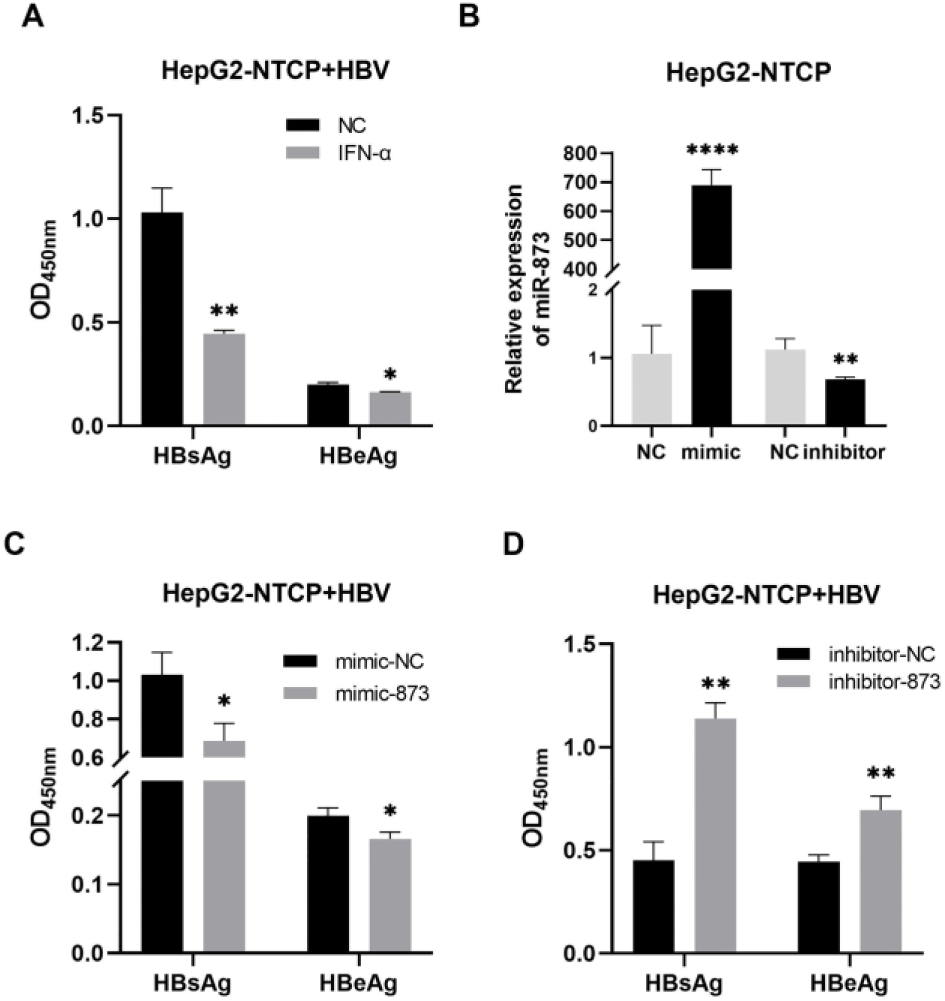
IFN-α and miR-873 suppresses HBV expression. (A) HBsAg and HBeAg expression were detected in HepG2-NTCP cells 48 h after treatment with IFN-α. (B) Efficiency assay of miR-873 mimic and inhibitor. (C) HBsAg and HBeAg expression were detected in HepG2-NTCP cells after treatment with miR-873 inhibitor. (D) HBsAg and HBeAg expression were observed after mimic-873 transfection. *, *P<*0.05; **, *P<*0.01; ****, *P<*0.0001.

### 3.6. Effect of miR-873 on the inhibition of HBV expression by IFN-*α*

As shown in Figure 6A and 6B, pHBV1.1-transfected HepG2-NTCP cells were treated with 1000 U/ml IFN-α and 20 nM miR-873 inhibitor. Unsurprisingly, IFN-α treatment and miR-873 inhibitor significantly reduced the level of HBsAg and HBeAg (*P*<0.05 or 0.01). Notably, the inhibitory effect was greatly enhanced by promoting the IFN-α-induced miR-873 downregulation by inhibitor transfection (*P*<0.01 or 0.001). In addition, pHBV1.1-transfected HepG2-NTCP cells were treated with IFN-α and miR-873 mimic as shown in Figure 6C and 6D. In NC-treated cells, IFN-α treatment resulted in a decrease in HBsAg and HBeAg level, whereas in miR-873 mimic-treated cells, the inhibitory effect was supressed by blocking the IFN-α-induced miR-873 downregulation by mimic transfection (*P*<0.01).

**Figure 6.**
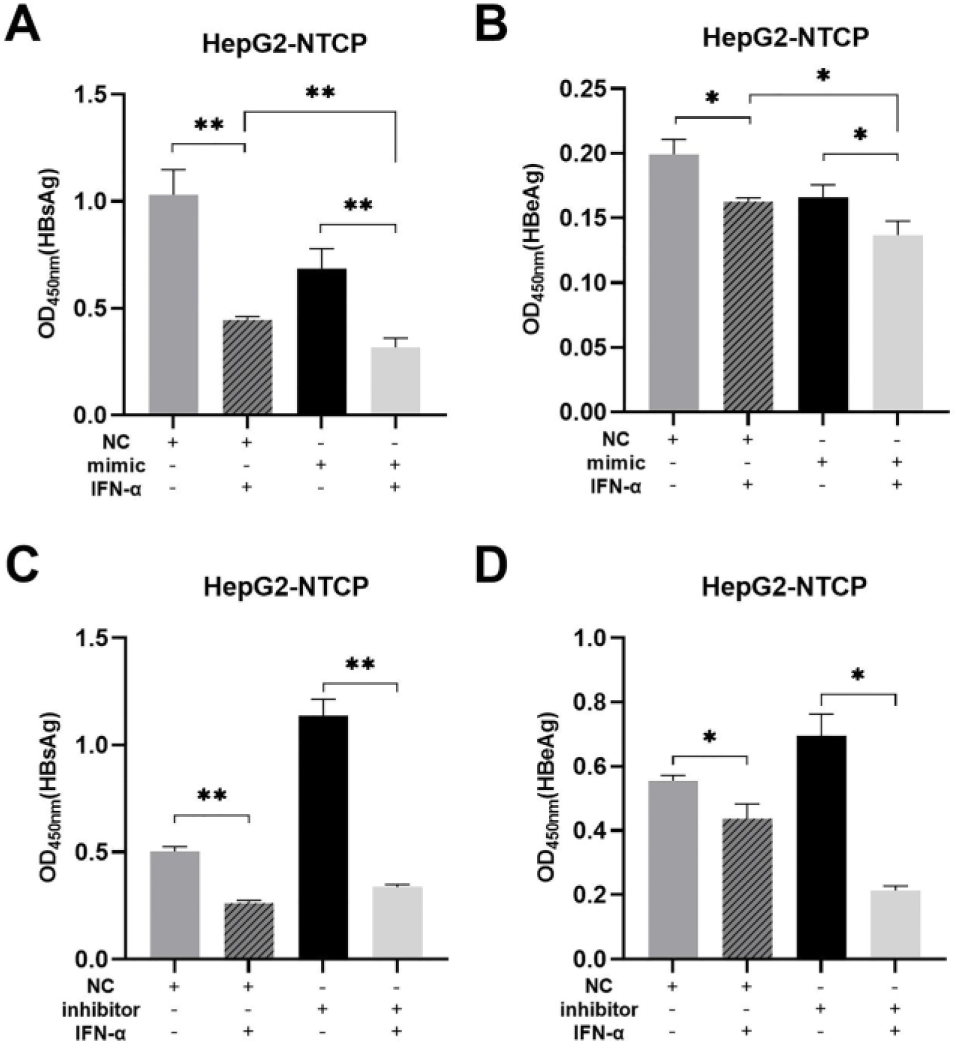
Effect of miR-873 on the down-regulation of HBV expression by IFN. (A) Effect of IFN treatment and inhibitor-873 on HBsAg level in HBV-infected cells. (B) Effect of IFN treatment and inhibitor-873 on HBeAg level in HBV-infected cells. (C) Effect of IFN treatment and mimic-873 on HBsAg level in HBV-infected cells. (D) Effect of IFN treatment and mimic-873 on HBeAg level in HBV-infected cells. *, P<0.05; **, P<0.01; ****, P<0.0001.

## 4. Discussion

CHB has brought a serious disease burden to humans. At present, a number of clinical studies have emphasized the importance of IFN in the treatment of hepatitis B and in sterilization therapeutic strategies[11, 20, 21]. Emerging data indicated that miRNAs were involved in host– virus interactions[22]. In this study, we explored the role of miR-873 in the development of chronic hepatitis B. We investigated the expression levels of miR-873, miR-200a, miR-30b, let-7g in PBMC of healthy volunteers and CHB patients and found that the expression levels of these miRNAs, especially miR-873, were upregulated. In addition, miR-873 inhibited the down-regulation of HBsAg and HBeAg by IFN in vitro. These results demonstrated that miR-873 plays a regulatory role in HBV biology, which provides a therapeutic strategy for chronic hepatitis B.

We identified differentially expressed miRNAs by machine learning algorithms. Machine learning models have been used to predict disease stages or therapeutic effects in the medical and health field for a long time. It has been reported to identify strong miRNAs associated with GC using Boruta machine learning variable selection approach[23]. In this study, miR-873, miR-200a, miR-30b, let-7g were identified as very strong candidates for IFN against HBV using the Random Forest algorithm. A prediction model based on these four core miRNAs showed 76% accuracy in predicting IFN responses.

Recent studies revealed that host miRNAs were associated with HBV replication[17, 24]. The molecular processes of HBV infection were a complex network, and miRNAs played different roles in this network. First, we determined the expression patterns of DEmiRNAs (miR-873, miR-200a, miR-30b and let-7g) in plasma in chronic hepatitis B, indicating that plasma DEmiRNAs mRNA expressions were enhanced in CHB. We further investigated the expression patterns of DEmiRNAs in the cells infected with HBV. Again, mRNA expressions of DEmiRNAs also significantly increased in HBV-infected cells. These findings revealed that miR-873 positively correlated with chronic hepatitis B which are in agreement with previous study in which miR-873 promoted viral replication. For instance, miR-873 promotes HIV-1 replication through HIV-1 gag, Pol, and p24 proteins[25]. Consistently, another report showed that miR-873 directly targeted p300 and p/CAF to affect chromatin remodeling and HTLV-1 replication[26].

Accumulating evidence demonstrated that dysregulation of host miRNAs affected type I IFN responses[27, 28]. After the expression patterns of miR-873 were identified, we then regulated the expressions of miR-873 and verified its effects on anti-HBV efficiency of IFN. High expressions of HBsAg indicate hepatitis B infection and reduction of HBsAg expression are considered an important goal in chronic hepatitis B treatment. HBeAg is located on the outer layer of the hepatitis B virus and its presence in patient plasma serves as an indicator of active viral replication. In the present study, we found that IFN induced down-regulation of HBsAg and HBeAg levels, whereas miR-873 mimic inhibited IFN-induced decrease in HBsAg and HBeAg. In contrast, miR-873 knockdown promoted IFN-induced downregulation of HBsAg and HBeAg levels. Furthermore, miR-873 affected immune function by altering the number of tumor-infiltrating immune cells or inhibiting macrophage apoptosis [29, 30]. These data revealed that miR-873 inhibit the down-regulation of HBsAg and HBeAg by IFN in vitro.

To date, IFN-α is a commonly administered treatment for individuals infected with HBV; however, this leads to a sustained antiviral response in only 25-40% of patients[31]. Given the difficulty in collecting a sufficient number of IFN-treated patients, we were unable to validate miR-873 expression in IFN-treated CHB patients. In further studies, more experiments will hopefully perform to validate the mechanism of action of miR-873 in HBV infection.

In conclusion, we provide evidence suggesting that miR-873 inhibition by IFN-α enhances the antiviral effects of IFN-α, as miR-873 is verified to promote HBV expression. Our work provides valuable insights to better understand the anti-HBV responses triggered by IFN-α and further provides potential strategies for enhancing the anti-HBV effects of IFN-α by inhibiting miR-873 expression.

## Data availability

All data generated and analyzed during the study are included in this published article,and detailed data are available at https://www.ncbi.nlm.nih.gov/geo/query/acc.cgi?acc=GSE29911.

---

**Supplementary Table 1.**
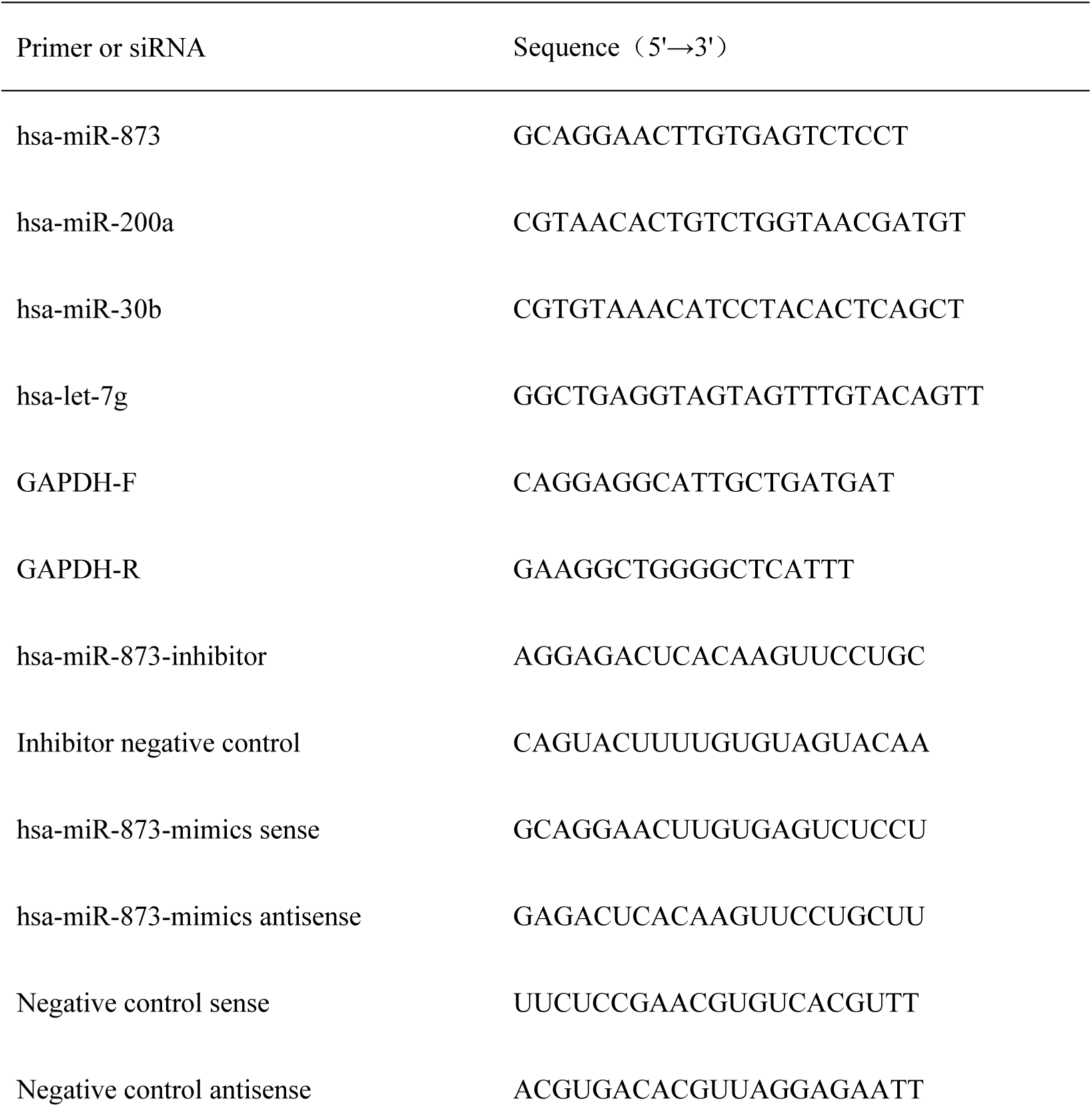
Sequences of primer and siRNA.

**Supplementary Table 2.** Baseline characteristics analysis of patients with chronic hepatitis B and normal controls.

| Characteristics | Total (N=36) | CHB (N=18) | NC (N=18) | P_Value |
| --- | --- | --- | --- | --- |
| Age | 33.50 (28.00, 15.50) | 35.00 (28.00, 49.50) | 32.00 (26.75, 44.50) | 0.424 |
| Male | 19(52.8) | 9 (50.0) | 10 (55.6) | 0.738 |
| ALT ( U/L) | 18.50 (13.25, 33.50) | 32.00 (19.00, 51.00) | 14.00(11.75, 17.25) | < **0.001** |
| AST (U/L) | 18.00 (15.25, 23.00) | 23.00(18.00,35.50 ) | 16.00(15.00,17.00 ) | < **0.001** |
| AST/ALT | 1.00 (0.77, 1.21) | 0.92(1.70,1.11 ) | 1.145(0.91,1.28 ) | **0.009** |
| ALP (U/L) | 72.00 (55.75, 92.00) | 76.00(62.50,97.25) | 69.50(47.75,88.25) | 0.265 |
| PAB(mg/L) | 223.5(195.50,<br>263.50) | 209.5(178.50,239.50) | 229(217.75,272.00) | **0.027** |
| TP (g/L) | 74.05 (70.65, 76.13) | 73.00(70.43,76.30) | 74.35(70.675,76.20) | 0.839 |
| ALB (g/L) | 45.45(43.10,47.45) | 45.90(44.68,48.20) | 44.95(42.78,47.13) | 0.293 |
| GLB(g/L) | 27.60 (25.23, 29.63) | 26.60(23.83,29.78) | 28.45(26.30,30.08) | 0.192 |
| A/G | 1.66 (1.44, 1.91) | 1.73(1.43,2.03) | 1.64(1.47,1.71) | 0.181 |
| TBIL (umol/L) | 11.30 (9.90, 16.63) | 10.60(7.43,13.10) | 14.40(10.30,17.95) | **0.068** |
| DBIL (umol/L) | 4.30 (2.98, 6.20) | 3.10(2.40,4.40) | 5.50(4.28,6.53) | **0.002** |
| IBIL (umol/L) | 7.55 (5.75, 10.93) | 7.40(4.40,8.58) | 8.90(6.05,11.33) | 0.203 |

ALT, alanine aminotransferase; AST, aspartate aminotransferase; ALP, alkaline phosphatase; PAB, prealbumin; TP, total protein; ALB, albumin; GLB, globulin; A/G, white globule ratio; TBIL, total bilirubin; DBIL, direct bilirubin; IBIL, indirect bilirubin.

**Supplementary Figure 1.**
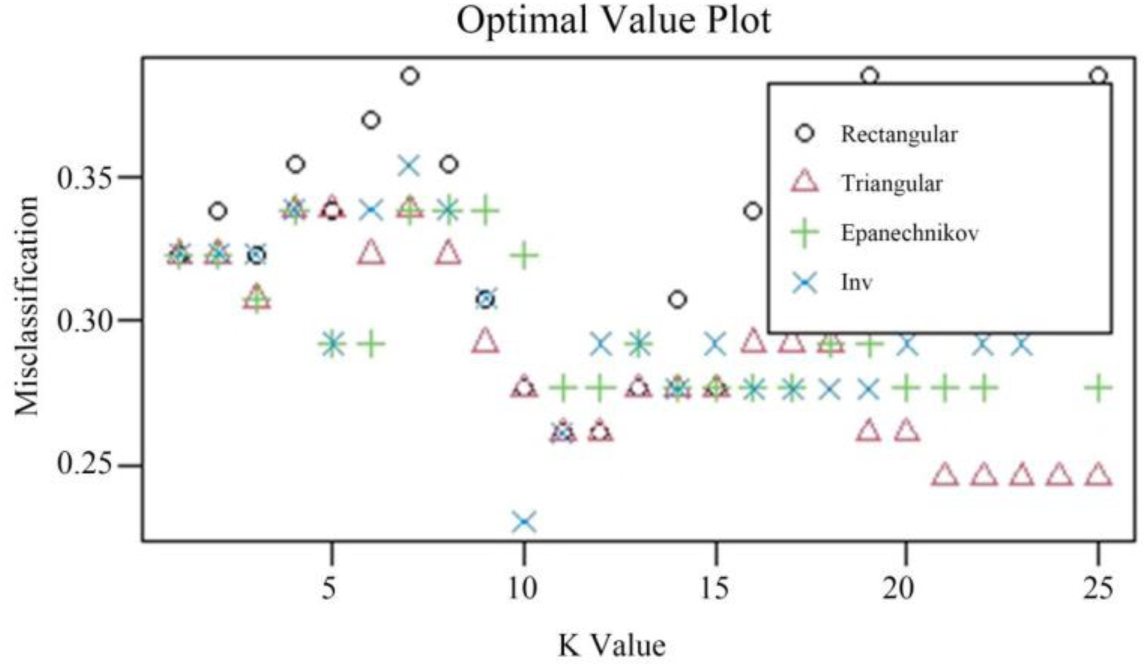
The prediction model constructed based on the KNN algorithm.

